# An RNA editing/binding-independent gene regulatory mechanism of ADARs and its clinical implication in cancer

**DOI:** 10.1101/062778

**Authors:** Lihua Qi, Yangyang Song, Tim Hon Man Chan, Henry Yang, Chi Ho Lin, Daryl Jin Tai Tay, HuiQi Hong, Jaymie Siqi Lin, Vanessa Hui En Ng, Julien Jean Pierre Maury, Daniel G. Tenen, Leilei Chen

## Abstract

Adenosine-to-inosine (A-to-I) editing, catalysed by Adenosine DeAminases acting on double-stranded RNA (dsRNA) (ADAR), occurs predominantly in the 3’ untranslated regions (3’UTRs). Here we uncover an unanticipated link between ADARs (ADAR1 and ADAR2) and the expression of target genes undergoing extensive 3’UTR editing. Using *METTL7A* (Methyltransferase Like 7A), a novel tumor suppressor as an exemplary target gene, we demonstrate that its expression could be repressed by ADARs beyond their RNA editing and dsRNA binding functions. ADARs interact with Dicer to augment the processing of pre-miR-27a to mature miR-27a. Consequently, mature miR-27a targets the *METTL7A* 3’UTR to repress its expression level. In sum, our study unveils that the extensive 3’UTR editing is merely a footprint of ADAR binding, and is dispensable for the regulation of at least a subset of target genes. Instead, ADARs contribute to cancer progression by regulating cancer-related gene expression through their non-canonical functions independent of RNA editing and dsRNA binding. The functional significance of ADARs is much more diverse than previously appreciated and this gene regulatory function of ADARs is most likely to be of higher importance than the best-studied editing function. This novel non-editing side of ADARs opens another door to target cancer. This study is timely and represents a major break-through in the field of ADAR gene regulation and cancer biology.

---

Adenosine DeAminases acting on dsRNA (ADAR) are highly conserved family of enzymes catalysing adenosine to inosine deamination (A-to-I editing) (Bass and Weintraub 1988; Wagner et al. 1989). There are 3 ADAR proteins (ADAR1, ADAR2 and ADAR3) in humans which all share a common modular structure characterized by 2 to 3 N-terminal dsRNA binding domains (dsRBDs) and a conserved C-terminal catalytic deaminase domain (Nishikura 2010; Qi et al. 2014). Being the best-studied function associated with ADAR1 and ADAR2 (ADARs), A-to-I RNA editing contributes to multi-level gene regulation depending on where it occurs. ADAR3, which has no documented deaminase activity, is only expressed in central nervous system (Melcher et al. 1996). In coding regions, A-to-I RNA editing can lead to a codon change and the consequent alterations of protein-coding sequences since inosine is interpreted by the ribosome as guanosine (Nishikura 2010). The differential editing frequencies of these recoding sites are found to impact on human diseases such as neurological diseases and cancer (Martinez et al. 2008; Chen et al. 2013; Galeano et al. 2013; Chan et al. 2014; Han et al. 2014; Paz-Yaacov et al. 2015; Villa et al. 2015). In non-coding regions, the vast majority of A-to-I RNA editing sites are in repetitive *Alu* elements embedded in 3’ untranslated regions (3’UTRs) (Farajollahi and Maas 2010) and have unknown functional relevance. Previously described fates of mRNAs undergoing extensive A-to-I editing at their 3’UTRs are via RNA editing-dependent mechanisms including nuclear retention, nuclease-mediated degradation, and alteration of microRNA (miRNA) targeting (Zhang and Carmichael 2001; Scadden 2005; Kawahara et al. 2007b; Morita et al. 2013), thereby influencing the expression of target genes.

ADARs have been found to be critical for normal development through and/or beyond A-to-I editing in different genetically modified animal models. Notably, the early post-natal lethality of the *Adar2−/−* mouse could be rescued by the ectopic expression of edited glutamate receptor subunit B (GluR-B), suggesting the editing activity of ADAR2 is essential for normal mouse development (Higuchi et al. 2000). However, whether ADAR1 editing activity is similarly responsible for the embryonic lethality of *Adarl−/−* mouse remains unclear, as the major editing targets of ADAR1 which are of high biological importance remain to be identified (Wang et al. 2000; Hartner et al. 2004). Furthermore, the primary microRNA (pri-miRNA) cleavage by Drosha/DGCR8 complex was also found to be inhibited by ADARs independent of their editing activities in both cell culture and the *Drosophila* models (Heale et al. 2009). Combining these editing-independent observations with the fact that humans have an editing-incompetent ADAR3 (Chen et al. 2000), it is most likely that ADARs can exert important functions as RNA binding proteins or through the formation of binding complexes with other proteins, beyond functioning as editing enzymes *per se*.

Hepatocellular carcinoma (HCC) is the most common type of liver malignancy, and accounts for approximately 700,000 cancer associated deaths per year worldwide (Ferlay et al. 2010). As reported previously, the deregulated RNA editing enzyme ADARs, as reflected by overexpression of ADAR1 and down-regulation of ADAR2, occurs in more than 50% of patients with HCC (Chan et al. 2014), rendering a disrupted RNA editing balance in both coding and non-coding regions (Chen et al. 2013; Chan et al. 2014). Moreover, the recurrent editing of *AZIN1* (antizyme inhibitor 1) which converts the codon 367 from serine to glycine has been demonstrated to predispose HCC development (Chen et al. 2013). However, most of A-to-I RNA editing occurs in the non-coding regions, and is particularly enriched in 3’UTRs (Peng et al. 2012). The contributions of 3’UTR editing by ADARs to cancer development have not yet been illustrated. Moreover, whether major regulatory mechanisms of ADARs on the expression of target genes with promiscuously edited 3’ UTRs are independent or dependent of their RNA editing capability, remain to be further explored.

To this end, we carried out the first systematic analysis of A-to-I editing events within 3’UTRs using our previously published RNA-Sequencing (RNA-Seq) datasets of 3 matched pairs of primary hepatocellular carcinoma (HCC) tumors and their adjacent non-tumor (NT) liver specimens (Chen et al. 2013; Chan et al. 2014), followed by the evaluation of a direct link between RNA editing at 3’UTRs and the expression of target transcripts. Surprisingly, a majority of target pre-mRNA transcripts with extensive editing at their 3’UTRs were found to be regulated by ADARs independent of their deaminase and dsRNA binding functions, providing new insights that the multiple A-to-I editing at 3’UTRs might be merely a footprint of ADAR binding, which is dispensable for the regulation of target gene expression. As an exemplary target gene with multiple editing sites at its 3’UTR, *METTL7A* could be regulated through a non-canonical regulatory mechanism of ADARs in which ADARs act as a modulator of the RNAi machinery rather than RNA editing. We also found that the downregulation of METTL7A by ADARs has biological implication in cancer development.

## Results

### Regulation by ADARs on the expression of target genes undergoing extensive A-To-I editing at 3’UTR sequences

Genes with extensive 3’UTR editing were selected based on our previously published RNA-sequencing (RNA-Seq) datasets(Chan et al. 2014), which have been generated from 3 paired primary HCC tumors and their adjacent NT liver specimens (Supplemental Data, Supplemental Fig. S1). To investigate if there exists any functional interaction between ADARs and the edited 3’UTRs, we constructed a 3’UTR-luciferase reporter system (Luc-3’UTRs) containing full-length 3’UTR sequences of 6 selected genes (*CCNYL1, TNFAIP8L1, MDM2, METTL7A, MTDH* and *RBBP9*) (Supplemental Data, Supplemental Fig. S2, Supplemental Table S1). Genetic structures of these 6 genes are illustrated in Fig. 1A. Co-transfection of Luc-3’UTRs together with the *ADAR1*-p110 (the isoform responsible for pre-mRNA editing(Farajollahi and Maas 2010)) or *ADAR2* expression constructs into HEK293T cells demonstrated that both *ADAR1* and *ADAR2* had suppressive effects on the luciferase activities linked to 3’UTRs of *TNFAIP8L1* and *METTL7A* gene, and the luciferase activities associated with *CCNYL1, MDM2* or *RBBP9* 3’ UTRs were only inhibited upon *ADAR2* overexpression; however, neither *ADAR1* nor *ADAR2* demonstrated any effect on *MTDH* 3’UTR (Fig. 1B, Supplemental Fig. S3).

**Figure 1.**
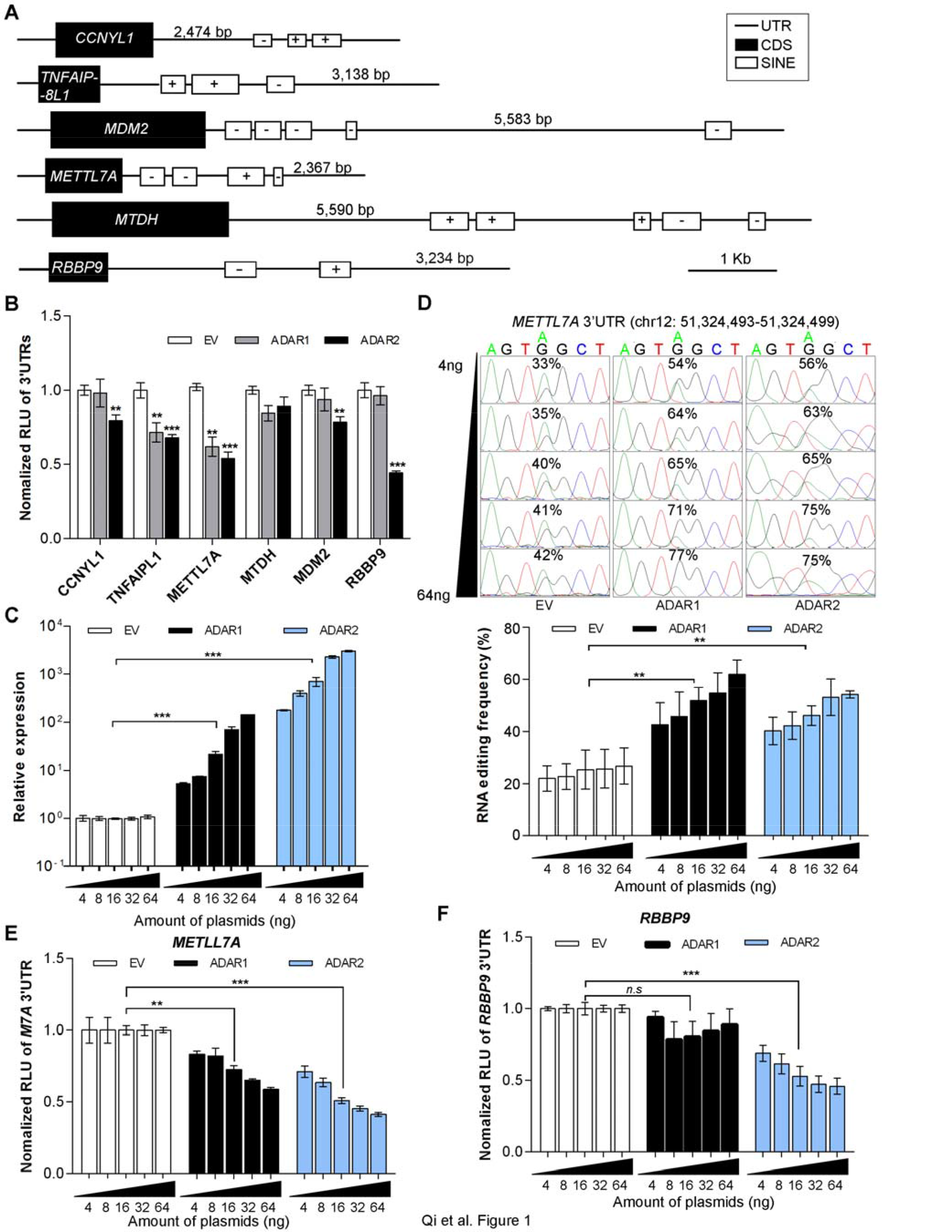
Regulatory effects of ADARs on target genes undergoing extensive 3’UTR editing. (*A*) Genetic structures of 6 selected genes. Black boxes represent the coding sequences (CDS), open boxes represent short interspersed elements (SINEs), and lines indicate untranslated regions (UTRs), using the RepeatMasker track of the UCSC genome browser(Kent et al. 2002). “+” and “−” signs indicate the strand specificity (sense or antisense. respectively) of SINEs. (*B*) Luciferase activities of pmirGLO-3’UTRs for the indicated genes were measured at 48 hours post co-transfection with the indicated expression constructs into HEK293T cells. RLU, relative luminescence unit. EV, empty vector (*C*) Quantitative real-time PCR (qRT-PCR) analyses of ADARs expression 48 hours post co-transfection of pmirGLO-METTL7A-3’UTR construct with increasing amounts of *ADARs* expression constructs into HEK293T cells. (*D*) Sequence chromatograms of one editing site in *METTL7A* 3’UTR in response to the co-transfection as described in (*C*). From top to bottom. the amount of *ADARs* expression construct was gradually increased from 4 ng to 64 ng. Percentages denote the editing frequencies of the corresponding editing sites (top panel). The bar chart represents the average editing frequency of 10 editing sites in *METTL7A* 3’UTR (identified by our RNA-Seq(Chan et al. 2014)) in each group of cells (bottom panel). (*E, F*), Luciferase activity of pmirGLO-*METTL7A*-3’UTR (*E*) or pmirGLO-*RBBP9*-3’UTR (*F*) measured 48 hours post co-transfection with increasing amounts of ADAR expression constructs into HEK293T cells. as described in (*A*). Data are presented as the mean ±s.e.m. of 6 replicates from a single experiment and representative of 3 independent experiments (*B-F*). Statistical significance was determined by unpaired. two-tailed Student’s *t*test, (*B-F*). (**, *P*< 0.01; ***, *P*<0.001; *n.s,* not significant).

Given the significantly reduced luciferase activities by *ADAR1* and/or *ADAR2, METTL7A* and *RBBP9* were selected for further investigation whether they are bona fide editing targets and more importantly, the suppressive effects of ADARs on their expression. In the light of the fact that a double stranded RNA (dsRNA) structure is essential for ADARs binding and A-to-I RNA editing (Farajollahi and Maas 2010), 3’UTR sequences of both *METTL7A* and *RBBP9*are predicted to form long dsRNA secondary structures using *CentroidFold* (Sato et al. 2009) (Supplemental Fig. S4). As expected, in HEK293T cells transfected with increasing amounts of either *ADAR1* or *ADAR2* expression construct, a dose-dependent reduction in luciferase activity of pmirGLO-METTL7A-3’UTR correlated negatively with the increase in the average editing frequency of 10 editing sites in the *METTL7A* 3’UTR (Fig. 1C-E; Supplemental Fig. S5A). As for *RBBP9,* we did observe a negative correlation between the luciferase activity of pmirGLO-RBBP9-3’UTR and the expression level of *ADAR2* but not with *ADAR1*, in a dose-dependent manner (Fig. 1F). Unexpectedly, none of 6 edited sites within the *RBBP9* 3’UTR demonstrated any increase in editing levels (Supplemental Fig. S5A, B). Altogether, we have shown that the expression of *METTL7A* and *RBBP9* could be regulated by ADAR1 and/or ADAR2; however whether this regulation is through an RNA editing-dependent or independent mechanism requires further investigation. Although we observed the significantly reduced 3’UTR-associated luciferase expression of *CCNYL1* and *TNFAIP8L1* upon *ADAR1* or/and *ADAR2* overexpression (Fig. 1B), there was no obvious change in the editing frequencies of their 3’UTR editing sites (Supplemental Fig. S5C). All these data suggests ADARs are most likely to regulate the expression of target genes undergoing extensive 3’UTR editing beyond their catalytic functions.

### RNA editing/dsRNA binding-independent suppression of ADARs on target genes

To this end, we generated different ADARs mutants devoid of either the enzymatic activity (DeAD mutants)(Lai et al. 1995) or the dsRNA binding capability (EAA mutants)(Valente and Nishikura 2007), which are both required for A-to-I RNA editing (Supplemental Methods, Supplemental Fig. S6). Editing frequencies of 10 editing sites within the *METTL7A* 3’UTR were examined in cells transfected with either wild-type or mutant form of ADARs (Fig. 2A, B). Consistent with a previous report (Valente and Nishikura 2007), ADARs DeAD mutants functioned as dominant negative forms, as they could compete for homodimerization with endogenous wild-type ADARs, rendering the wild-type partners inactive. As for ADARs EAA mutants, there was no significant difference in the basal editing level of *METTL7A* 3’UTR when compared to the control (ADAR1 EAA: *P* = 0.14; ADAR2 EAA: *P* = 0.47) (Fig. 2A, B), which might be attributed to the loss of dimerization ability of EAA mutants (Poulsen et al. 2006). Consistent with the data described above (Fig. 1B), luciferase activities of pmirGLO-*METTL7A*-3’UTR and pmirGLO-*TNFAZP8L1*-3’UTR could be suppressed by both wild-type and mutant forms of ADARs (Fig. 2C, E). Luciferase activities linked to 3’UTRs of *CCNYL1* and *RBBP9* were only suppressed by different forms of *ADAR2* in HEK293T cells (Fig. 2D, F). Similar effects of different ADAR forms on luciferase activities of pmirGLO-METTL7A-3’UTR and pmirGLO-RBBP9-3’UTR were also observed in the HCC cell line SNU-398 (Supplemental Fig. S7). All these data strongly suggests a non-canonical regulatory function of ADARs on gene expression through 3’UTRs independent of RNA editing and dsRNA binding.

**Figure 2.**
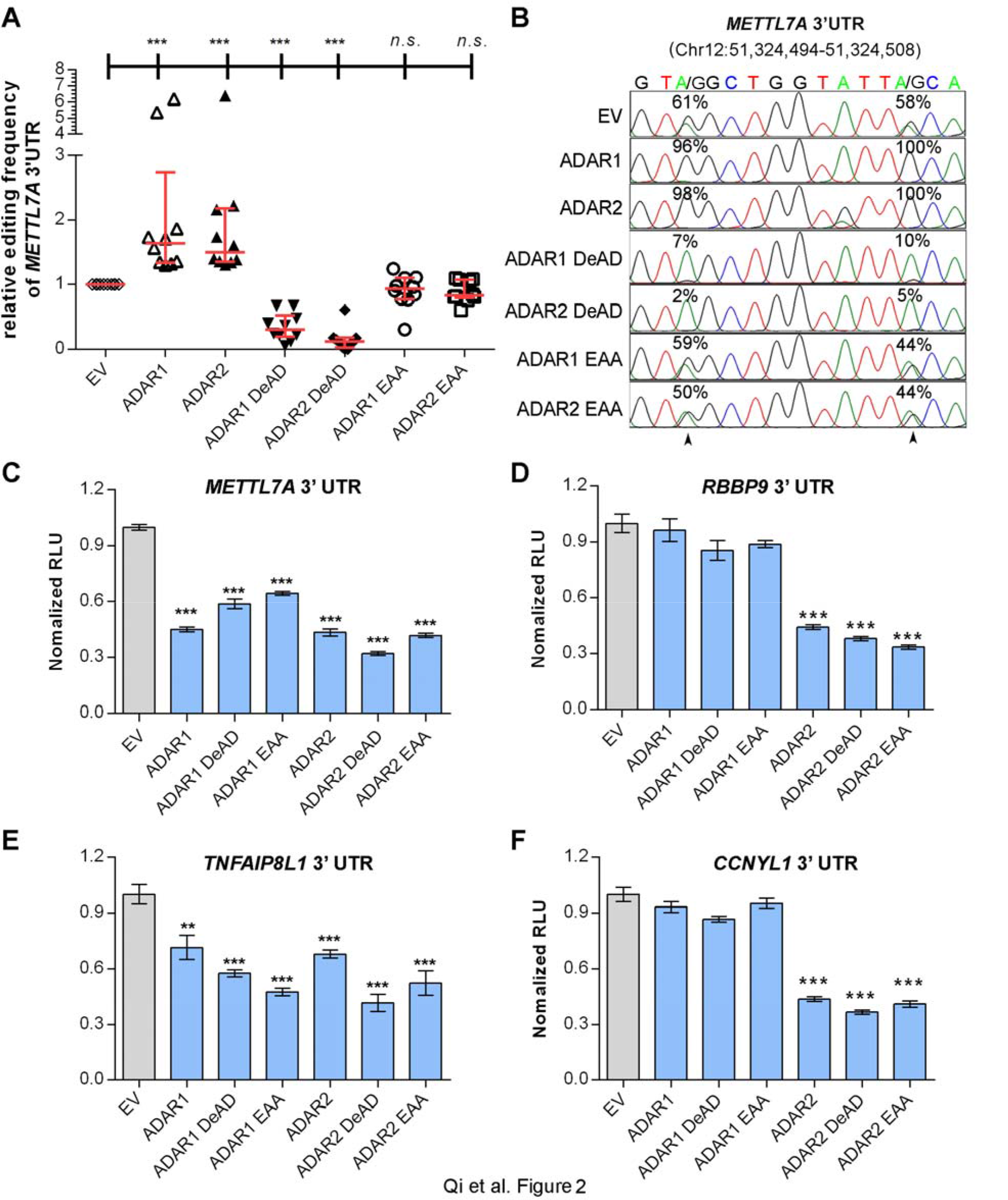
RNA editing/dsRNA binding-independent suppression of target genes by ADARs. (*A*) Scatter plots showing the relative editing frequencies of 10 editing sites at *METTL7A* 3’UTR in response to overexpression of different ADARs (wild-type: ADAR1 or ADAR2; DeAD mutant: ADAR1 DeAD or ADAR2 DeAD; dsRNA-binding mutant: ADAR1 EAA or ADAR2 EAA) in HEK293T cells. The data are presented with median (horizontal line) and interquartile range (error bar) for each group (Mann-Whitney U test; ***, *P*< 0.001; *n.s*, not significant). (*B*) Sequence chromatograms of 2 editing sites within *METTL7A* 3’UTR. Percentages denote the editing frequencies of the corresponding editing sites indicated by arrowheads. (*C-F*) Luciferase activity of pmirGLO-*METTL7A*-3’UTR (*C*), pmirGLO-*RBBP9*-3’UTR (*D*), pmirGLO-*TNFAIP8L1*-3’UTR (*E*), or pmirGLO-*CCNYL1*-3’UTR (*F*) 48 hours post co-transfection with the control empty vector (EV) or different ADARs constructs (wild-type and mutants) in HEK293T cells. Data are presented as the mean ± s.e.m. of 6 replicates from a single experiment and representative of 3 independent experiments (**, *P* < 0.01; ***, *P* < 0.001). Statistical significance was determined by unpaired. twotailed Student’s *t* test (*C-F*).

### Suppression by ADARs on *METTL7A* expression in HCC cells and primary tumors

To demonstrate that our observations have biological implications in cancer development, beyond exogenous luciferase reporter assays, the *METTL7A* gene was selected for further investigation of ADARs-mediated effects on endogenous gene expression. Due to the neutral effects of EAA mutants on A-to-I editing and its loss of dsRNA binding capability, EAA mutants of ADARs were included in the remainder of this study. Both wild-type and EAA mutants inhibited endogenous METTL7A expression at both mRNA and protein levels in the HCC cell line Huh-7, which has the highest endogenous METTL7A expression among 9 HCC cell lines (Fig. 3A, B; Supplemental Fig. S8A). Conversely, specific shRNAs against *ADAR1* or *ADAR2* increased METTL7A expression in the HCC cell line SMMC7721, expressing relatively high levels of ADARs among 9 cell lines (Fig. 3C; Supplemental Fig. S8B, C). Altogether, ADARs could exert RNA editing and dsRNA binding-independent suppression on *METTL7A* expression in cancer cells.

**Figure 3.**
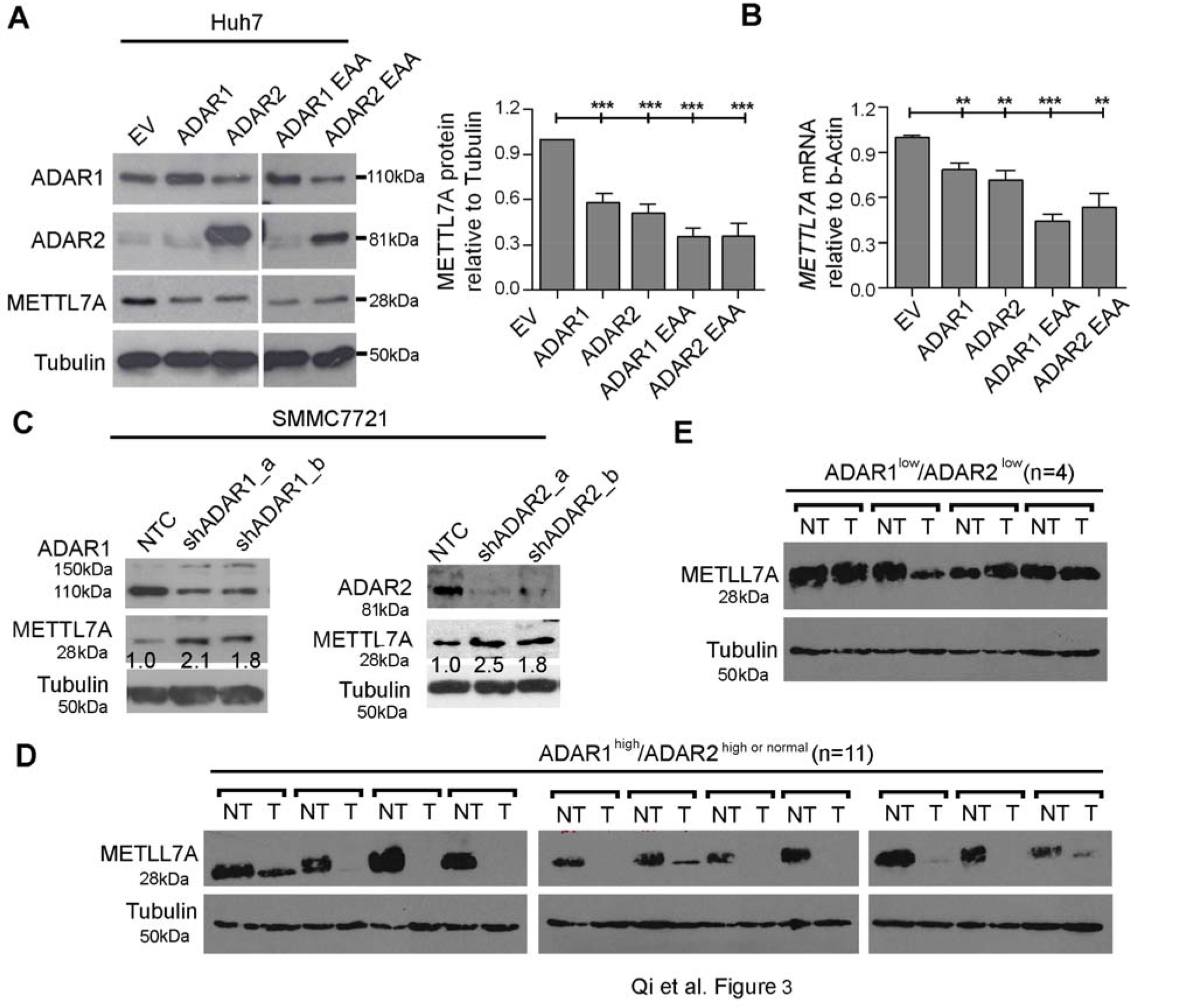
RNA editing/dsRNA binding-independent suppression by ADARs on *METTL7A* expression in HCC cells and primary tumors. (*A*) Western blot analyses of the indicated proteins in Huh-7 cells 72 hours after transfection with the wild-type or EAA mutants of ADARs (left panel). The bar chart represents the normalized densitometry unit of METTL7A protein on Western blot (right panel). (*B*) qRT-PCR analysis of *METTL7A* mRNA level measured in the same samples as described in (*A*). (*C*) Western blot analyses of the indicated proteins SMMC7721 cells 72 hours post transfection with shRNAs against either *ADAR1* or *ADAR2*. Value indicates the normalized densitometry unit of METTL7A protein on Western blot. (*D-E*) Western blot analysis of METTL7A protein in 2 groups of HCC cases including ADAR1^high^/ADAR2^high or normal^ (*D*) and ADAR1^low^/ADAR2^low^ (*E*). Tubulin was the loading control. Data are presented as the mean ± s.e.m. of 6 replicates from a single experiment and representative of 3 independent experiments (*A, B*).Statistical significance was determined by unpaired. two-tailed Student’s *t* test (*A* and *B*). (**, *P* < 0.01; ***, *P* < 0.001).Qi et al.

To study if this non-canonical regulatory mechanism is of high biological importance during cancer development, we went on to examine ADARs-mediated suppression on METTL7A expression in 15 paired primary HCC tumor and their matched NT liver specimens. Patients with HCC were divided into 2 groups: ADAR1^high^/ADAR2^high or normal^ (n=11) and ADAR1 ^low^/ADAR2^low^ (n=4) (Supplemental Methods, Supplemental Fig. S9). When compared to their matched NT specimens, all of ADAR1^hlgh^/ADAR2^hlgh or normal^ HCCs demonstrated either lower or absent expression of METTL7A (Fig. 3D); while 3 out of 4 (75%) ADAR1^low^/ADAR2^low^ HCCs demonstrated an increase or no change in METTL7A expression (Fig. 3E). This suggests the ADARs-mediated suppression may be a major mechanism of *METTL7A* downregulation in HCCs.

### MiR-27a as a mediator between ADARs and *METTL7A*

In light of the fact that 3’UTR is the major miRNA targeting region, miRNAs are most likely to mediate this non-canonical regulation of ADARs on *METTL7A* expression. Editing-independent effects of ADARs on RNA interference (RNAi) and miRNA processing, which were first delineated in *Drosophila* (Heale et al. 2009), have been recently observed in mammalian systems (Ota et al. 2013).

In order to identify miRNAs targeting *METTL7A*, the *METTL7A* 3’UTR sequence was subjected to miRNA prediction using miRWalk across multiple algorithms (miRanda, miRDB, miRWalk, RNA22 and Targetscan) (Dweep et al. 2011). Only miRNAs that can be predicted by at least 2 algorithms were selected for further investigation. As seen in Fig. 4A, the introduction of miR-27a into HEK293T cells significantly repressed the luciferase activity of pmirGLO-*METTL7A*-3’UTR, indicating that miR-27a might truly target the *METTL7A* 3’UTR. Further, the luciferase activity of pmirGLO-*METTL7A*-3’UTR was found to be repressed by miR-27a, in a dose-dependent manner (Fig. 4B). Subsequently, the pmirGLO-*METTL7A*-3’UTR mutant was constructed by mutating the seed region (indicated by box; Fig. 4C) in the pmirGLO-METTL7A-3’UTR construct. Transfection of the pmirGLO-*METTL7A*-3’UTR mutant into HEK293T cells completely abolished the inhibition by miR-27a of the luciferase activity (Fig. 4C), indicating a direct interaction between miR-27a and the *METTL7A* 3’UTR. Moreover, the expression of endogenous METTL7A protein was also found to be inhibited by miR-27a in Huh-7 cells (Fig. 4D).

**Figure 4.**
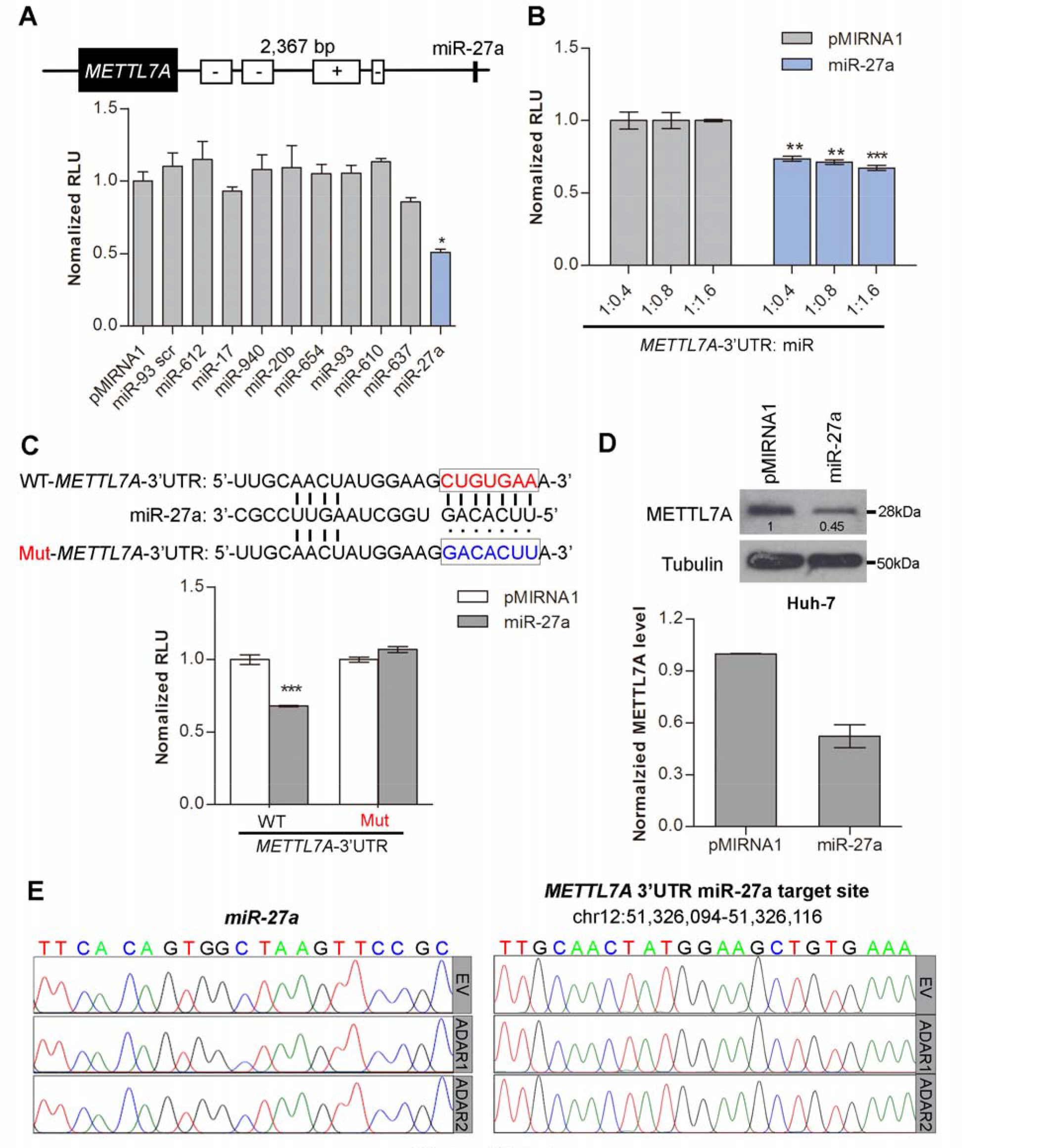
MiR-27a as a mediator between ADARs and *METTL7A*. (*A*) Schematic diagram demonstrates the predicted miR-27a targeting region on the *METTL7A* 3’UTR. The bar chart presents the luciferase activity of pmirGLO-METTL7 *A*-3’UTR 48 hours after co-transfection with the indicated miRNAs at a mass ratio of 1:9 in HEK293T cells. pMIRNA1, empty vector; miR-93 scr, scrambled control (scrambled mature miR-93 seed sequence). (*B*) Luciferase activity of pmirGLO-*METTL7A*-3’UTR measured 48 hours post co-transfection with the increasing amount of miR-27a in HEK293T cells (**, *P*< 0.01; ***, *P*<0.001). (*C*) Schematic diagram showing the mutations being introduced to the predicted miR-27a target site of the *METTL7A* 3’UTR (top panel). WT, wild-type; Mut, mutant. The bar chart presents the luciferase activity associated with the wild-type or mutant pmirGLO-*METTL7A* 3’UTR 48 hours after co-transfection with miR-27a in HEK293T cells (bottom panel). (*D*) Western blot analysis of METTL7A protein in Huh-7 cells stably overexpressing miR-27a. Value indicates the normalized densitometry unit of METTL7A protein on Western blot. The bar chart presents the densitometry unit of METTL7A from 3 independent experiments. Tubulin was the loading control. (*E*) Sequence chromatograms of miR-27a and the predicted miR-27a target sequence within the *METTL7A* 3’UTR, in Huh-7 cells stably overexpressing miR-27a as described in (*A*). Data are presented as the mean ± s.e.m. of 6 replicates from a single experiment and representative of 3 independent experiments(*A-C*). Statistical significance was determined by unpaired. two-tailed Student’s *t* test (*A-D*). (*, *P* < 0.05; **, *P* < 0.01; ***, *P* < 0.001).

Importantly, we further confirmed that there was no A-to-I editing occurring in miR-27a and its targeted *METTL7A* 3’UTR sequences, ruling out the possibility that the impact of ADARs on *METTL7A* expression is via the enhancement of miR-27a:*METTL7A* mRNA interaction arising from the A-to-I editing of miR-27a and/or its target *METTL7A* 3’UTR sequence (Fig. 4E).

### ADARs interact with dicer to promote miR-27a expression independent of RNA editing/dsRNA binding activities

If miR-27a is a key mediator for ADARs-mediated suppression of *METTL7A*, both the wild-type and EAA mutants of ADARs are predicted to upregulate miR-27a. Indeed, mature miR-27a was found to be upregulated upon overexpression of either wild-type or EAA mutants of ADARs in Huh-7 cells (Fig. 5A). We went on to study which stage of miR-27a biosynthesis could be targeted by ADARs. There was no obvious increase in the expression of miR-27a precursors, pri-miR-27a, and pre-miR-27a, upon the overexpression of either wild-type or EAA mutant (Fig. 5B, C). Further, Northern blot analysis for miR-27a confirmed that the introduction of both wild-type and EAA forms of ADARs could process pre-miR-27a to mature miR-27a more efficiently than the control (Fig. 5D). All these observations indicated that processing of pre-miR-27a to mature miR-27a is likely to be regulated by ADARs independent of RNA editing and dsRNA binding.

**Figure 5.**
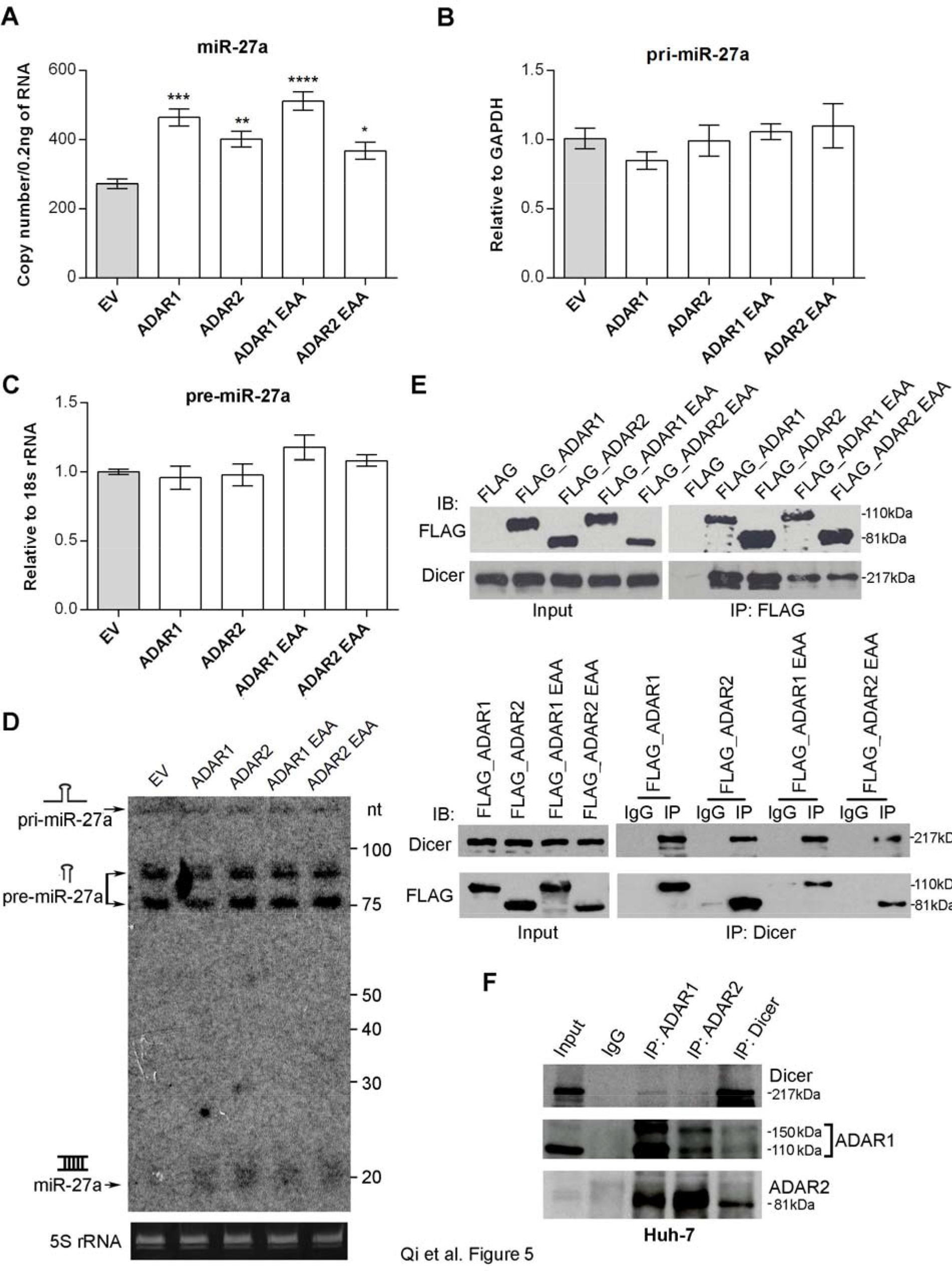
ADARs interact with Dicer to promote miR-27a expression independent of RNA editing/dsRNA binding activities. (*A-C*) The qRT-PCR measurement of mature miR-27a (*A*), pri-miR-27a (*B*), and pre-miR-27a (*C*) in Huh-7 cells upon overexpression of empty vector (EV) or different ADARs expression constructs including the wild-type and EAA mutants. Data are presented as the mean ± s.e.m. of triplicates from a single experiment and representative of 3 independent experiments (\**P* < 0.05; \*\**P* < 0.01; \*\*\**P* < 0.001). Statistical significance was determined by unpaired. two-tailed Student’s *t* test. (*D*) Northern blot analysis of mature miR-27a, pri-miR-27a, and pre-miR-27a in same samples as in A. 5S rRNA bands stained with ethidium bromide are presented as a loading control. (*E*) Reciprocal immunoprecipitation (IP) of ADARs and Dicer was performed in HEK293T cells overexpressing FLAG only or FLAG-tagged wild-type and mutant form of ADARs. using anti-FLAG antibody conjugated magnetic beads (IP: FLAG; top panel), or Dicer-specific antibody (IP: Dicer; bottom panel) or mouse IgG (IP: IgG; bottom panel). Western blot analyses of the indicated proteins in FLAG. Dicer or mouse IgG-IPed products were conducted. (*F*) Reciprocal IP of endogenous ADARs and Dicer was conducted in Huh-7 cells using an ADAR1, ADAR2 or Dicer-specific antibody (IP: ADAR1, ADAR2 or Dicer) or mouse IgG (IP: IgG). Western blot analyses of the indicated proteins in ADAR1, ADAR2, Dicer or mouse IgG-IPed products were conducted. Input indicates 5% of whole-cell lysates from HEK293T cells transfected with the indicated constructs (*E*) or Huh-7 cells(*F*).

Processing of pre-miRNAs to miRNAs is catalysed by Dicer in complex with TRBP (Tar RNA binding protein)(Winter et al. 2009). Also, it has been recently reported that ADAR1 promote pre-miRNA processing via binding to Dicer (Ota et al. 2013). Therefore, we hypothesized that both ADARs promote miR-27a processing and expression through interacting with Dicer. As expected, endogenous Dicer was detected in both FLAG-tagged wild-type and EAA mutant pull-down products (Fig. 5E). Reciprocally, FLAG-tagged wild-type ADARs and EAA mutants could also be detected in anti-Dicer immunoprecipitates (Fig. 5E). More importantly, the reciprocal pull-down assays indicated endogenous Dicer could interact with both ADAR1 and ADAR2 in Huh-7 cells (Fig. 5F). The observed interaction between ADAR1 and ADAR2, known to form heterodimer in human cells (Chilibeck et al. 2006; Cenci et al. 2008), serves as positive controls. Taken together, ADARs are found to augment the processing of pre-miR-27a to mature miR-27a via binding to Dicer, in turn targeting the *METTL7A* 3’UTR and decreasing *METTL7A* expression.

### *METTL7A* as a novel tumor suppressor in HCC

Having identified a non-canonical function of ADARs beyond their catalytic and dsRNA-binding capabilities, we sought to demonstrate that the altered expressions of target genes has a biological impact on cancer development. *METTL7A* expression was analysed using RNA-Seq datasets from the Cancer Genome Atlas (TCGA) project (Weinstein et al. 2013) and a microarray gene expression dataset from the NCBI Gene Expression Omnibus (GEO) database (GEO542361) (Villa et al. 2015). Both datasets confirmed significantly decreased expression of *METTL7A* in HCC tumors compared to adjacent NT tissues, and also predicted shorter overall survival for patients with HCC (Fig. 6A, B).

**Figure 6.**
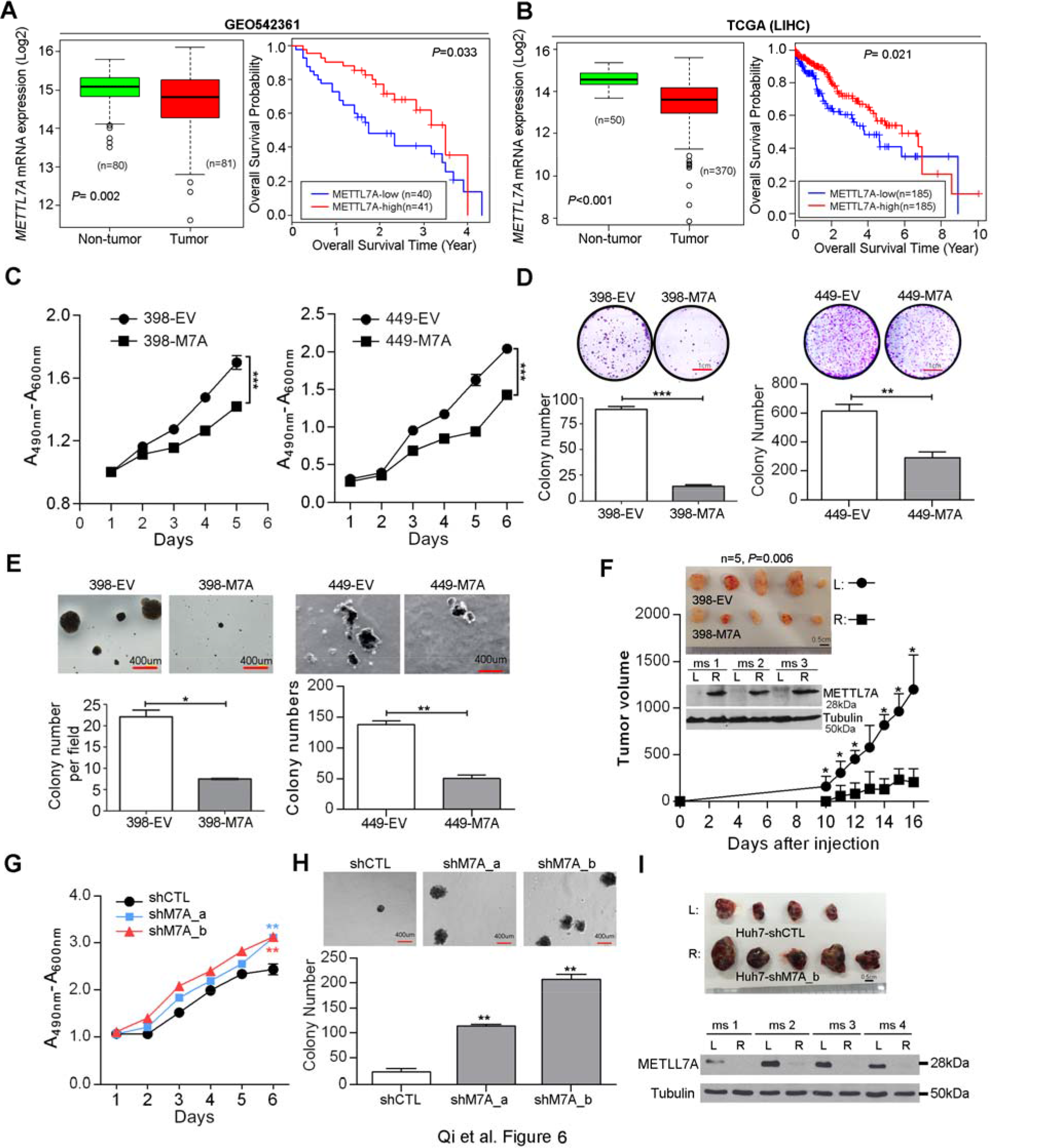
*METTL7A* is a novel tumor suppressor in HCC. (*A*, *B*) Box plots showing *METTL7A* expression in HCC and matched NT liver specimens. as retrieved from the GEO54236 microarray gene expression datasets in (*A*), from TCGA (Liver Hepatocellular carcinoma. LIHC) RNA-Seq datasets in (B). The data are presented as box plots with median (horizontal line), 25-75% (box) and 5-95% (error bar) percentiles for each group and the open dots indicate the outliers (Mann-Whitney U test). Kaplan-Meier plots for the overall survival rate of patients with HCC in the group with high(METTL7A-high: n=41 for GEO54236; n=185 for LIHC) or low *METTL7A* expression (METTL7A-low: n=40 for GEO54236; n=185 for LIHC) which was categorized based on the median expression of *METTL7A* in HCCs. The *P* value shown was calculated by log rank test. (*C*) 2,3-bis-(2-methoxy-4-nitro-5-sulfophenyl)-2H-tetrazolium-5-carboxanilide (XTT) assay showing growth rates of the indicated stable cell lines. A_490nm_ and A_600nm_, absorbance at 490 nm and 600 nm, respectively. (***, *P* < 0.001). (*D*) Quantification of foci formation induced by the indicated stable cell lines (**, *P* < 0.01; ***, *P*< 0.001). Scale bar. 1cm. (*E*) Quantification of soft agar colonies induced stable cell lines (*, *P* < 0.05; **, *P* < 0.01). Scale bar. 400 μm. (*F*) Tumors derived from the indicated cell lines 3 weeks after subcutaneous injection (n=5 mice per group. top panel). Scale bar. 0.5 cm. Western blot analysis of METTL7A protein in tumors developed in 3 representative mice at end point (middle panel). Growth curves of tumors derived from the indicated stable cell lines over the observation period (bottom panel). Data are presented as the mean ± s.e.m. *, *P* < 0.05, determined by unpaired. twotailed Student’s *t* test. (*G*) XTT assay showing growth rates of the stably knockdown cell lines. A_490nm_ and A_600nm_, absorbance at 490 nm and 600 nm, respectively. (**, *P* < 0.01). (*H*) Quantification of soft agar colonies induced stably knockdown cell lines (**, *P*± 0.01). Scale bar. 400 μm. (*I*) Top panel shows the xenograft tumors derived from the indicated stable cell lines 4 weeks post subcutaneous injection (n=5 mice per group). Scale bar. 0.5 cm. Western blot analysis of METTL7A protein in tumors developed in 4 representative mice at end point (bottom panel). Tubulin was the loading control. All data are shown as the mean ± s.e.m. of triplicate wells with the same experiment and representative of 3 independent experiments (*C, D, E, G, H*),and statistical significance was determined by unpaired. two-tailed Student’s *t* test (*C-H*).

To further investigate the role of METLL7A in HCC development, 3 HCC cell lines (SNU-398, SNU-449, and Huh-7) were selected for establishing overexpression and knockdown cell models and the subsequent functional studies (Fig. S10A, B). As detected by cell culture assays predictive of tumorigenicity, overexpression of METTL7A in both SNU-449 and SNU-398 cells (398-M7A and 449-M7A) significantly reduced the cell viability, frequency of focus formation, and the number of colonies formed in soft agar when compared to cells transduced with empty vector lentiviruses (398-EV & 449-EV) (Fig. 6C-E). Moreover, xenograft tumors derived from 398-M7A cells, with overexpression of METTL7A, grew much slower than tumors derived from 398-EV cells during a 3-week observation period (Fig. 6F). Conversely, specific shRNAs-mediated silencing of *METTL7A* augmented the tumorigenicity of Huh-7 cells, as indicated by the increased cell viability and higher frequency of colony formation in soft agar (Fig. 6G, H; Fig. S10C). As seen in Fig. 6I, xenograft tumors derived from stably knockdown cells (Huh7-shM7A_b) grew much more aggressively than tumors derived from control cells (Huh7-shCTL), as a consequence of the persistent silencing of *METTL7A* during a 4-week observation period (Fig. 6I). All these data suggests that METTL7A downregulation, possibly mainly due to the RNA editing/binding-independent suppression by ADARs, is closely linked to HCC development.

## Discussion

Extensive A-to-I editing at 3’UTR sequences has been associated with different fates of the edited pre-mRNA transcripts. Due to the distinct base pairing preference between adenosine and inosine, RNA editing at 3’UTRs can regulate the stability of target transcripts by creating or eliminating miRNA targeting sites (Borchert et al. 2009; Wang et al. 2013). Alternatively, the promiscuously edited 3’UTRs can recruit inosine-specific nucleases, e.g. the RNA-induced silencing complex (RISC) subunit Tudor-SN and human endonuclease V, leading to mRNA degradation (Scadden 2005; Morita et al. 2013). Certain transcripts with edited 3’UTRs were shown to be retained in the nucleus by an inosine-specific nuclear protein complex composed of P54^nrb^, splicing factor PSF, and matrin-3 (refs (Prasanth et al. 2005; Nishikura 2010)). All of the above reported fates of inosine-containing RNAs highlighted the importance of inosine in either changing the base-pairing properties or recruiting inosine-specific protein complexes, thereby affecting the translational efficiency of the edited mRNAs. In contrast, a previous study in *C. elegans* reported no significant difference in expression of mRNAs with structured and edited 3’UTRs across different *adr* strains (Hundley et al. 2008). Furthermore, mRNAs with edited 3’UTRs have been demonstrated to associate with translating polyribosomes in both *C. elegans* and human HeLa cells, indicating that not all the edited mRNAs are retained in the nucleus (Hundley et al. 2008). Thus, a relevant question is whether RNA editing is necessary to induce the observed effect of ADARs on 3’UTRs?

Surprisingly, using different ADAR mutants which are devoid of either RNA editing or dsRNA binding capability, we found the promiscuous A-to-I editing within 3’UTRs of transcripts is most likely to be a footprint of ADARs binding, which is not required for the manipulation of target gene expression, at least for a subset of targets. Relatively few protein variants produced by RNA editing have been demonstrated to be causative of biological phenotypes observed in animal models with genetically modified ADARs suggesting that editing-independent functions of ADARs might also be of importance in the observed phenotypes (Nie et al. 2005). Furthermore, a recent study suggested that the ADAR1 binding to non-*Alu* regions could affect 3’UTR usage through competing with 3’UTR-binding factors, such as cleavage and polyadenylation-relevant proteins (Bahn et al. 2015), thereby influencing the expression of target genes. More importantly, the impact of ADAR1 on 3’UTR usage is only dependent on RNA editing for certain 3’UTRs, but others could be affected by ADAR1 in an editing-independent manner. In this study, 5 out of the 6 selected genes carrying multiple editing sites at 3’UTRs were found to be regulated by ADAR1 and/or ADAR2 beyond their deaminase and dsRNA binding functions. Therefore, the functional significance of ADARs is much more diverse than previously appreciated and this gene regulatory function of ADARs may be of higher importance than the better-studied editing function.

Until now, gene regulatory mechanisms of ADARs have not been clearly understood. Given the fact that ADAR-mediated A-to-I RNA editing competes for shared dsRNA substrates with RNAi machinery, interactions between ADARs and RNAi pathway have gained intensive attention during the past decade (Nishikura 2006). Effects of hyper-editing in protecting dsRNA from entering RNAi were first confirmed by biochemical assays, and further observed in *C. elegans* with *adr-1/adr-2* double mutant models (Scadden and Smith 2001; Tonkin and Bass 2003). Editing of primary miRNAs has been reported to inhibit miRNA biosynthesis at multiple stages, e.g. Drosha and Dicer cleavage, the loading to RISC, or redirecting silencing targets (Yang et al. 2006; Kawahara et al. 2007a; Kawahara et al. 2007b; Kawahara et al. 2008; Iizasa et al. 2010). In contrast to antagonistic effects of ADARs on the RNAi pathway, a recent study on the impact of *ADAR1* on pri-miRNA processing in the nucleus suggested that ADAR1 might enhance pri-mRNA processing via its interaction with the pri-mRNAs prior to Drosha/DGCR8 binding, which may not rely on RNA editing (Bahn et al. 2015). This finding complements the previous report that the interaction between ADAR1 and RISC component proteins revealed an editing-independent agonistic effect of ADAR1 on miRNA processing, by forming a complex with Dicer in the cytoplasm (Ota et al. 2013). Here, our mechanistic studies focused on *METTL7A*, an exemplary target gene with multiple editing sites in its 3’UTR, indicated that the interaction of ADARs with Dicer augments the processing from pre-miR-27a to mature miR-27a and subsequent miR-27a targeting of the *METTL7A* 3’UTR, leading to reduced expression of *METTL7A*. Thus, our study represents the first report that both *ADAR1* and *ADAR2* can augment miRNA processing from pre-miRNAs to mature miRNAs through interacting with *Dicer*, independent of RNA editing and dsRNA binding. It is likely that a large amount of genes are regulated by ADARs via non-canonical editing/dsRNA binding-independent mechanism(s), such as *CCNYL1, MDM2, TNFAIP8L1*, and *RBBP9*, which have been identified in this study. However, for other target genes, the precise regulatory mechanism(s) of ADARs will need to be examined on a case-by-case basis.

The next question is whether this mechanism may have important functional relevance in cancer development. As an exemplary target gene undergoing 3’UTR editing, *METTL7A* was identified to be a tumor suppressor in HCC. Moreover, the survival analysis of 2 individual HCC cohorts indicated the tumoral downregulation of *METTL7A* predicted poor prognosis of patients with HCC. Unexpectedly, even with multiple editing sites in the 3’UTR, the expression of METTL7A was found to be regulated by ADARs through miRNA expression modulation in an RNA editing and RNA binding-independent manner. Given the widespread effects of miRNA targeting, *METTL7A* will likely not be the only target influenced by ADARs under this mechanism. Recent studies reported the expression of miRNAs is globally inhibited in *adar1^−/−^* mouse embryos, in turn altering the expression of miRNA target genes (Ota et al. 2013). Also, in human U87MG cells, a gene otology (GO) analysis of target genes of ADAR1-affected miRNA yields a number of genes functionally enriched in pathways related to cell proliferation, growth, apoptosis and cellular response to stimuli or DNA damage (Bahn et al. 2015).

Our study revealed that the functional essentiality/importance of ADARs is likely to stem from its involvement in processes other than RNA editing and dsRNA binding alone, at least for a subset of target genes relevant to cancer development. As we and other have demonstrated, the enhancement of miRNA processing via the interaction between ADARs and Dicer (Ota et al. 2013) or Drosha (Bahn et al. 2015) may function as a major mechanism responsible for gene regulatory function of ADARs beyond their deaminase and RNA binding activities. Disruption of the interaction between ADARs and Dicer or Drosha might represent a targetable approach for rescuing expression of tumor suppressors such as *METTL7A* as a novel therapeutic modality in HCC and other cancers.

## Methods

### Cell lines

SNU-398, SNU-449, and HEK293T cell lines were purchased from the American Type Culture Collection (ATCC). Huh-7 was purchased from Japanese Collection of Research Bioresources Cell Bank (JCRB). SMMC-7721 was purchased from Thermo Fisher Scientific, MA, USA. SNU-398 and SNU-449 cells were cultured in Roswell Park Memorial Institute (RPMI) medium (Gibco BRL, Grand Island, NY, USA) supplemented with 10% fetal bovine serum (FBS) (Gibco BRL). Huh-7, SMMC-7721, and HEK293T cells were maintained in Dulbecco’s Modified Eagle’s medium (DMEM) supplemented with 10% FBS. All cell lines used in this study were regularly authenticated by morphological observation and tested for mycoplasma contamination (MycoAlert, Lonza Rockland, Rockland, ME). The cells were incubated at 37 °C in a humidified incubator containing 5% CO_2_.

### Clinical samples

Total of 15 paired human primary HCC and adjacent NT liver tissues that were surgical removed and snap frozen in liquid nitrogen were obtained from the tissue repository of National University Hospital (NUH), Singapore. All patients gave written informed consent for the use of their clinical specimens for medical research. All samples used in this study were approved by the Institutional Review Board of National University of Singapore (NUS-IRB) (Reference Code: B-14-239E).

### Publically available databases

Two publically available datasets for survival data analysis of patients with HCC with respect to *METTL7A* expression: RNA-Seq datasets (including 370 HCC tumors and 50 adjacent NT liver samples) from TCGA (LIHC)(Weinstein et al. 2013) (https://tcga-data.nci.nih.gov/tcga/) and microarray gene expression datasets (including 81 HCC tumors and 80 adjacent NT liver samples) from GEO54236(Villa et al. 2015) (http://www.ncbi.nlm.nih.gov/gds). Prior to the survival analysis, raw RNA-Seq counts were normalized using the total numbers of mappable reads across all samples, while the microarray data normalization was performed using the Cross-Correlation method (Chua et al. 2006). Normalized and log2 transformed *METTL7A* expression data were subjected to the survival analysis using Kaplan-Meier plots and log-rank tests. The median tumoral *METTL7A* expression was used to define METTL7A-high and METTL7A-low groups of patients with HCC for both datasets. The *METTL7A* expression in tumors and matched NT tissues were compared using the Mann-Whitney U test.

### Statistical analysis

The SPSS statistical package for Windows, version 16 (SPSS), and Microsoft Excel (Excel in Microsoft Office 2013 for Windows) were used for data analysis. The expression of METTL7A in tumors and matched NT tissues were compared using the Mann-Whitney U test. The relative editing frequencies of 10 representative editing sites at *METTL7A* 3’UTR in response to overexpression of the control or different ADARs in HEK293cells were compared using the Mann-Whitney U test. Kaplan-Meier plots and log-rank tests were used for overall survival analysis. The unpaired, two-tailed Student’s *t* test was used to compare the number of foci, colony formation, tumor volume and relative expressions of target genes, the luciferase activities between any two preselected groups. *P* < 0.05 was considered statistically significant.

## Author Contributions

L. Q., Y. S., L. C. and D.G.T. initiated the study. L. Q. wrote the manuscript with inputs from T.H.M.C., H.Q., J.J.P.M., D.G.T, L. C., Y.S. and J.S.L. L.Q., Y.S. and L. C. designed the experiments. L.Q., Y.S. and L.C. performed all experiments with assistance from T.H.M.C., H.Q., L. C., J.S.L., D.J.T.T. and V.N. C.H.L. performed all bioinformatics analyses of RNA Sequencing datasets. H.Y., L. Q., Y. S. and L. C. contributed to data acquisition, analysis, and interpretation. L. C. and D.G.T. supervised the project.

## Acknowledgments

We thank and acknowledge the patients for tumor tissue donation and Dr. J. Yan (Cancer Science Institute of Singapore, National University of Singapore, Singapore) for providing the pMIRNA1 lenti-miR expression vector. This work was supported by the National Research Foundation Singapore, and the Singapore Ministry of Education under its Research Centres of Excellence initiative, NMRC Clinician Scientist-Individual Research Grant New Investigator Grant (CS-IRG NIG, grant number: NMRC/CNIG/1117/2014), NMRC Clinician Scientist-Individual Research Grant (CS-IRG, grant number: NMRC/CIRG/1412/2014), NUS Young Investigator Award (NUS YIA, grant number: NUSYIA_FY14_P22), Yong Siew Yoon Research Grant (Tier 2; National University Cancer Institute, Singapore), NUS Start-up fund (Ref. No: NUHSRO/2015/095/SU/01) and the Singapore Ministry of Health’s National Medical Research Council under its Singapore Translational Research (STaR) Investigator Award. This research is also supported by the RNA Biology Center at the Cancer Science Institute of Singapore, NUS, as part of funding under the Singapore Ministry of Education’s Tier 3 grants (MOE2014-T3-1-006).

## References

Bahn JH., Ahn J., Lin X., Zhang Q., Lee JH., Civelek M., Xiao X. 2015. Genomic analysis of ADAR1 binding and its involvement in multiple RNA processing pathways. Nat Commun 6: 6355.

Bass BL., Weintraub H. 1988. An unwinding activity that covalently modifies its double-stranded RNA substrate. Cell 55: 1089–1098.

Borchert GM., Gilmore BL., Spengler RM., Xing Y., Lanier W., Bhattacharya D., Davidson BL. 2009. Adenosine deamination in human transcripts generates novel microRNA binding sites. Hum Mol Genet 18: 4801–4807.

Cenci C., Barzotti R., Galeano F., Corbelli S., Rota R., Massimi L., Di Rocco C., O’Connell MA., Gallo A. 2008. Down-regulation of RNA editing in pediatric astrocytomas: ADAR2 editing activity inhibits cell migration and proliferation. J Biol Chem 283: 7251–7260.

Chan TH., Lin CH., Qi L., Fei J., Li Y., Yong KJ., Liu M., Song Y., Chow RK., Ng VH et al. 2014. A disrupted RNA editing balance mediated by ADARs (Adenosine DeAminases that act on RNA) in human hepatocellular carcinoma. Gut 63: 832–843.

Chen CX., Cho DS., Wang Q., Lai F., Carter KC., Nishikura K. 2000. A third member of the RNA-specific adenosine deaminase gene family. ADAR3, contains both single-and double-stranded RNA binding domains. RNA 6: 755–767.

Chen L., Li Y., Lin CH., Chan TH., Chow RK., Song Y., Liu M., Yuan YF., Fu L., Kong KL et al. 2013. Recoding RNA editing of AZIN1 predisposes to hepatocellular carcinoma. Nat Med 19: 209–216.

Chilibeck KA., Wu T., Liang C., Schellenberg MJ., Gesner EM., Lynch JM., MacMillan AM. 2006. FRET analysis of in vivo dimerization by RNA-editing enzymes. J Biol Chem 281: 16530–16535.

Chua SW., Vijayakumar P., Nissom PM., Yam CY., Wong VV., Yang H. 2006. A novel normalization method for effective removal of systematic variation in microarray data. Nucleic Acids Res 34: e38.

Dweep H., Sticht C., Pandey P., Gretz N. 2011. miRWalk–database: prediction of possible miRNA binding sites by “walking” the genes of three genomes. J Biomed Inform 44: 839–847.

Farajollahi S., Maas S. 2010. Molecular diversity through RNA editing: a balancing act. Trends in genetics: TIG 26: 221–230.

Ferlay J., Shin HR., Bray F., Forman D., Mathers C., Parkin DM. 2010. Estimates of worldwide burden of cancer in 2008: GLOBOCAN 2008. Int J Cancer 127: 2893–2917.

Galeano F., Rossetti C., Tomaselli S., Cifaldi L., Lezzerini M., Pezzullo M., Boldrini R., Massimi L., Di Rocco CM., Locatelli F et al. 2013. ADAR2-editing activity inhibits glioblastoma growth through the modulation of the CDC14B/Skp2/p21/p27 axis. Oncogene 32: 998–1009.

Han SW., Kim HP., Shin JY., Jeong EG., Lee WC., Kim KY., Park SY., Lee DW., Won JK., Jeong SY et al. 2014. RNA editing in RHOQ promotes invasion potential in colorectal cancer. JExp Med 211: 613–621.

Hartner JC., Schmittwolf C., Kispert A., Muller AM., Higuchi M., Seeburg PH. 2004. Liver disintegration in the mouse embryo caused by deficiency in the RNA-editing enzyme ADAR1. J Biol Chem 279: 4894–4902.

Heale BS., Keegan LP., McGurk L., Michlewski G., Brindle J., Stanton CM., Caceres JF., O’Connell MA. 2009. Editing independent effects of ADARs on the miRNA/siRNA pathways. EMBO J 28: 3145–3156.

Higuchi M., Maas S., Single FN., Hartner J., Rozov A., Burnashev N., Feldmeyer D., Sprengel R., Seeburg PH. 2000. Point mutation in an AMPA receptor gene rescues lethality in mice deficient in the RNA-editing enzyme ADAR2. Nature 406: 78–81.

Hundley HA., Krauchuk AA., Bass BL. 2008. C. elegans and H. sapiens mRNAs with edited 3’ UTRs are present on polysomes. RNA 14: 2050–2060.

Iizasa H., Wulff BE., Alla NR., Maragkakis M., Megraw M., Hatzigeorgiou A., Iwakiri D., Takada K., Wiedmer A., Showe L et al. 2010. Editing of Epstein-Barr virus-encoded BART6 microRNAs controls their dicer targeting and consequently affects viral latency. J Biol Chem 285: 33358–33370.

Kawahara Y., Megraw M., Kreider E., Iizasa H., Valente L., Hatzigeorgiou AG., Nishikura K. 2008. Frequency and fate of microRNA editing in human brain. Nucleic Acids Res 36: 5270–5280.

Kawahara Y., Zinshteyn B., Chendrimada TP., Shiekhattar R., Nishikura K. 2007a. RNA editing of the microRNA-151 precursor blocks cleavage by the Dicer-TRBP complex. EMBO Rep 8: 763–769.

Kawahara Y., Zinshteyn B., Sethupathy P., Iizasa H., Hatzigeorgiou AG., Nishikura K. 2007b. Redirection of silencing targets by adenosine-to-inosine editing of miRNAs. Science 315: 1137–1140.

Kent WJ., Sugnet CW., Furey TS., Roskin KM., Pringle TH., Zahler AM., Haussler D. 2002. The human genome browser at UCSC. Genome research 12: 996–1006.

Lai F., Drakas R., Nishikura K. 1995. Mutagenic analysis of double-stranded RNA adenosine deaminase. a candidate enzyme for RNA editing of glutamate-gated ion channel transcripts. J Biol Chem 270: 17098–17105.

Martinez HD., Jasavala RJ., Hinkson I., Fitzgerald LD., Trimmer JS., Kung HJ., Wright ME. 2008. RNA editing of androgen receptor gene transcripts in prostate cancer cells. J Biol Chem 283: 29938–29949.

Melcher T., Maas S., Herb A., Sprengel R., Higuchi M., Seeburg PH. 1996. RED2, a brain-specific member of the RNA-specific adenosine deaminase family. J Biol Chem 271: 31795–31798.

Morita Y., Shibutani T., Nakanishi N., Nishikura K., Iwai S., Kuraoka I. 2013. Human endonuclease V is a ribonuclease specific for inosine-containing RNA. Nat Commun 4: 2273.

Nie Y., Ding L., Kao PN., Braun R., Yang JH. 2005. ADAR1 interacts with NF90 through doublestranded RNA and regulates NF90-mediated gene expression independently of RNA editing. Mol Cell Biol 25: 6956–6963.

Nishikura K. 2006. Editor meets silencer: crosstalk between RNA editing and RNA interference. Nat Rev Mol Cell Biol 7: 919–931.

Nishikura K. 2010. Functions and regulation of RNA editing by ADAR deaminases. Annu Rev Biochem 79: 321–349.

Ota H., Sakurai M., Gupta R., Valente L., Wulff BE., Ariyoshi K., Iizasa H., Davuluri RV., Nishikura K. 2013. ADAR1 forms a complex with Dicer to promote microRNA processing and RNA-induced gene silencing. Cell 153: 575–589.

Paz-Yaacov N., Bazak L., Buchumenski I., Porath HT., Danan-Gotthold M., Knisbacher BA., Eisenberg E., Levanon EY. 2015. Elevated RNA Editing Activity Is a Major Contributor to Transcriptomic Diversity in Tumors. Cell reports 13: 267–276.

Peng Z., Cheng Y., Tan BC., Kang L., Tian Z., Zhu Y., Zhang W., Liang Y., Hu X., Tan X et al. 2012. Comprehensive analysis of RNA-Seq data reveals extensive RNA editing in a human transcriptome. Nat Biotechnol 30: 253–260.

Poulsen H., Jorgensen R., Heding A., Nielsen FC., Bonven B., Egebjerg J. 2006. Dimerization of ADAR2 is mediated by the double-stranded RNA binding domain. RNA 12: 1350–1360.

Prasanth KV., Prasanth SG., Xuan Z., Hearn S., Freier SM., Bennett CF., Zhang MQ., Spector DL. 2005. Regulating gene expression through RNA nuclear retention. Cell 123: 249–263.

Qi L., Chan TH., Tenen DG., Chen L. 2014. RNA editome imbalance in hepatocellular carcinoma. Cancer research 74: 1301–1306.

Sato K., Hamada M., Asai K., Mituyama T. 2009. CENTROIDFOLD: a web server for RNA secondary structure prediction. Nucleic Acids Res 37: W277–280.

Scadden AD. 2005. The RISC subunit Tudor-SN binds to hyper-edited double-stranded RNA and promotes its cleavage. Nat Struct Mol Biol 12: 489–496.

Scadden AD., Smith CW. 2001. RNAi is antagonized by A-->I hyper-editing. EMBO Rep 2: 1107–1111.

Tonkin LA., Bass BL. 2003. Mutations in RNAi rescue aberrant chemotaxis of ADAR mutants. Science 302: 1725.

Valente L., Nishikura K. 2007. RNA binding-independent dimerization of adenosine deaminases acting on RNA and dominant negative effects of nonfunctional subunits on dimer functions. J Biol Chem 282: 16054–16061.

Villa E., Critelli R., Lei B., Marzocchi G., Camma C., Giannelli G., Pontisso P., Cabibbo G., Enea M., Colopi S et al. 2015. Neoangiogenesis-related genes are hallmarks of fast-growing hepatocellular carcinomas and worst survival. Results from a prospective study. Gut doi:10.1136/gutjnl-2014-308483.

Wagner RW., Smith JE., Cooperman BS., Nishikura K. 1989. A double-stranded RNA unwinding activity introduces structural alterations by means of adenosine to inosine conversions in mammalian cells and Xenopus eggs. Proc Natl Acad Sci U S A 86: 2647–2651.

Wang Q., Hui H., Guo Z., Zhang W., Hu Y., He T., Tai Y., Peng P., Wang L. 2013. ADAR1 regulates ARHGAP26 gene expression through RNA editing by disrupting miR-30b-3p and miR-573 binding. RNA 19: 1525–1536.

Wang Q., Khillan J., Gadue P., Nishikura K. 2000. Requirement of the RNA editing deaminase ADAR1 gene for embryonic erythropoiesis. Science 290: 1765–1768.

Weinstein JN., Collisson EA., Mills GB., Shaw KRM., Ozenberger BA., Ellrott K., Shmulevich I., Sander C., Stuart JM., Network CGAR. 2013. The Cancer Genome Atlas Pan-Cancer analysis project. Nat Genet 45: 1113–1120.

Winter J., Jung S., Keller S., Gregory RI., Diederichs S. 2009. Many roads to maturity: microRNA biogenesis pathways and their regulation. Nat Cell Biol 11: 228–234.

Yang W., Chendrimada TP., Wang Q., Higuchi M., Seeburg PH., Shiekhattar R., Nishikura K. 2006. Modulation of microRNA processing and expression through RNA editing by ADAR deaminases. Nat Struct Mol Biol 13: 13–21.

Zhang Z., Carmichael GG. 2001. The fate of dsRNA in the nucleus: a p54(nrb)-containing complex mediates the nuclear retention of promiscuously A-to-I edited RNAs. Cell 106: 465–475.

